# Optimized fluorescent proteins for 4-color and photoconvertible live-cell imaging in *Neurospora crassa*

**DOI:** 10.1101/2022.09.12.507692

**Authors:** Ziyan Wang, Bradley M. Bartholomai, Jennifer J. Loros, Jay C. Dunlap

## Abstract

Fungal cells are quite unique among life in their organization and structure, and yet implementation of many tools recently developed for fluorescence imaging in animal systems and yeast has been slow in filamentous fungi. Here we present analysis of properties of fluorescent proteins in *Neurospora crassa* as well as describing genetic tools for the expression of these proteins that may be useful beyond cell biology applications. The efficacy of ten different fluorescent protein tags were compared in a constant context of genomic and intracellular location; six different promoters are described for the assessment of the fluorescent proteins and varying levels of expression, as well as a customizable bidirectional promoter system. We present an array of fluorescent proteins suitable for use across the visible light spectrum to allow for 4-color imaging, in addition to a photoconvertible fluorescent protein that enables a change in the color of a small subset of proteins in the cell. These tools build on the rich history of cell biology research in filamentous fungi and provide new tools to help expand research capabilities.

## 1. Introduction

The fungal cell is a marvel of evolutionary engineering. Hyphae, as we know them, are unique among life in their long tubular structure that is highly branched and interconnected. With many nuclei sharing a crowded cytoplasm that is racing through the interconnected network of hyphae we know as a mycelium, fungi elicit the question, “What is a cell?”. The microscopic study of filamentous fungi dates back to the late 17^th^ century, when Marcello Malpighi published the purported first hand drawn micrograph of hyphae (Malpighi, 1675-1679; Money, 2021). Today, filamentous fungi are the subject of a wide variety of microscopy based cellular investigations.

Fluorescence based tools are a cornerstone of contemporary light microscopy-based cell biology research. Since use of the *Aequorea victoria* green fluorescent protein was introduced in the early 1990s to tag individual proteins and visualize their subcellular localization, a vast toolbox of fluorescent proteins has been developed to visualize subcellular phenomena across the visible light spectrum (Chalfie et al., 1994; Rodriguez et al., 2017). By 2019, the number of reported fluorescent proteins had grown to such a large number that the community driven FPbase database was formed to help researchers compare properties of different proteins and choose the best tools for their needs (Lambert, 2019). However, the vast majority of these proteins have been optimized for animal systems for which the intracellular milieu and even amino acid codon preferences may be different from those in fungi; additionally, reported values for various protein characteristics including brightness, aggregation, pKa, etc. have been determined *in vitro* or are calculated from theoretical values (Campbell et al., 2020; Day and Davidson, 2009; Hirano et al., 2022). In fungi, these proteins can perform quite differently than how one might expect from the reported parameters.

While several publications have compared and optimized fluorescent proteins in yeast, few examples of such work can be found for filamentous fungi (Higuchi-Sanabria et al., 2016; Lee et al., 2013; Schuster et al., 2015). Unfortunately, tools that work well in yeast do not always perform similarly in filamentous fungi. Advances in fluorescence microscopy of yeast have also outpaced filamentous fungi in the use of multiple (up to four) different fluorescent proteins to visualize different proteins in the same living cell (Higuchi-Sanabria et al., 2016). To address this gap, we have built upon previous work and developed a set of tools to help expand the repertoire of fluorescent protein reporters in filamentous fungi, focusing in the present case on *Neurospora crassa*.

Since the first reported use of *A. victoria* GFP in *N. crassa* in 2001, the fluorescent toolbox in Neurospora has expanded modestly to include tdimer2, dsRed, YFP, and mCherry, with GFP and mCherry being most common (Bardiya et al., 2008; Freitag et al., 2001; Freitag and Selker, 2005; Verdín et al., 2009). The tools presented here were developed as part of our efforts to understand the spatiotemporal dynamics of the *Neurospora crassa* circadian clock, for which we had very specific needs that were not met by previously described reporters. For context, the circadian clock is a single step negative feedback loop in which a complex of proteins scaffolded by FRQ facilitates the phosphorylation of its transcriptional activators, WC-1 and WC-2 acting as the White Collar Complex of WCC, depressing its activity. Eventual phosphorylation of FRQ causes it to lose affinity for the WCC, allowing resynthesis of new FRQ (Diernfellner and Brunner, 2020; Dunlap and Loros, 2017; Larrondo et al., 2015). Because many of these proteins are expressed at a very low level, to ensure physiological relevance we needed very bright fluorescent proteins that would allow us to resolve signal from these proteins at their endogenous expression levels. We also needed to be able to image multiple proteins concurrently, visualize rapid localization changes, and express reporters and varying constitutive expression levels.

Here we present analysis of brightness and photobleaching properties for a number of fluorescent proteins in *N. crassa*, studies that revealed a green fluorescent protein that is significantly brighter than the most commonly used GFP in Neurospora. We also optimized a set of fluorescent proteins that can be used for simultaneous 4-color (blue, green, red, near infrared) imaging in living hyphae. In addition, we present an optimized photoconvertible protein that facilitates a color change from green to red after a pulse of violet light thereby allowing estimation of intracellular mixing rates. These tools can all be driven by a series of promoters we describe for varying constitutive expression levels of heterologous protein constructs or over/under expression. Through application of these tools we observed simultaneous bidirectional transport of separate nuclei through septal pores both antero- and retrograde to bulk flow as previously described (Mouriño-Pérez et al., 2016; Ramos-García et al., 2009). We were also able to track movements of nuclei from just one region of a hypha through implementation of a photoconvertible fluorescent protein and observed apparently stress-related ring-like and tubular septal structures not seen before in Neurospora. We offer these tools to the community with hopes of expanding the possibilities of investigating the fascinating cellular biology of filamentous fungi.

## 2. Results and Discussion

### 2.1 Optimization and *in vivo* evaluation of constitutively expressed fluorescent proteins in *N. crassa*

Experiments in circadian biology involve the study of proteins expressed in a circadian manner that are relatively low in abundance even at their peak. Cells must also be imaged over time periods relevant to circadian biology (≥ 24 h). Therefore, we sought out the brightest and most stable fluorescent proteins (FPs) that have been described in the literature. Unfortunately, the values provided for brightness, photobleaching, and other parameters that are presented in databases such as FPbase (Lambert, 2019) are either theoretical, measured *in vitro*, or most often based on values obtained from mammalian cells. To study fluorescent markers that could be used to examine the transcription-translation negative feedback loop that comprises the molecular clock, we required a system that could be used to systematically test fluorescent proteins as well as providing a versatile set of nuclear periphery markers. We decided to use the outer nuclear envelope nucleoporin SON-1 for this purpose.

SON-1(NCU04288) was previously identified as a homolog to SONA in *Aspergillus nidulans* (Roca et al., 2010). These proteins are homologous to the highly conserved eukaryotic RAE1/GLE2 proteins (NCBI HomoloGene:2676). In *Aspergillus*, it was shown to dissociate from the nuclear envelope during mitosis (De Souza et al., 2004); however, this was not observed in *Neurospora* (Roca et al., 2010). As a component of the nuclear pore complex (NPC), this protein interacts with Nup98 (human) on the cytoplasmic ring of the NPC (Von Appen and Beck, 2016), and has been shown to be involved in RNA nuclear export, septin organization, spindle assembly, and even nucleocytoplasmic shuttling of the core circadian CLOCK/BMAL complex in mammals (Kato et al., 2021; Murphy et al., 1996; Pritchard et al., 1999; Wong, 2010; Zander et al., 2017; Zheng et al., 2019). Because SON-1^GFP^ had been expressed in *Neurospora* previously, we used it as a nuclear envelope marker for our work. Additionally, because SON-1 does not leave the nucleus and nuclei are discrete and easy to observe, SON-1 provided a useful platform from which to examine and compare the efficacy of various candidate fluorescent proteins in vivo in Neurospora. Given this, our screening strategy was to develop a plasmid capable of (1) driving stable expression of *son-1* tagged with various fluorescent proteins (FP) and (2) integrating into a fixed neutral site in the genome.

In considering a promoter to drive *son-1^FP^* expression, we realized that the most commonly used promoter for such reporters was *ccg-1;* however, *ccg-1* is highly and conditionally regulated by light, oxygen levels, and nutrition as well as by the clock (Arpaia et al., 1995; Loros et al., 1989; McNally and Free, 1988), so we queried data from Hurley et al. (Hurley et al., 2015; Hurley et al., 2018) seeking a constitutive promoter with a level of expression allowing easy visualization of the proteins to be tested. From this analysis we chose NCU04502 which is constitutively transcribed under our conditions and encodes a small “hypothetical protein” that is constitutively translated. To develop the promoter of NCU04502 for use in screening diverse fluorescent proteins, we used SON-1 tagged with luciferase and targeted to the *csr-1* locus for its ease of initial screening, testing several lengths of nucleotides upstream of the NCU04502 transcriptional start site (Supplemental Figure 1A) for expression by measuring bioluminescence of transformants. Subsequent western blot analysis using a luciferase antibody (Santa Cruz Biotechnology Cat. #sc-74548) confirmed high luciferase expression from the constructs containing 600 to 1000 bp of upstream DNA, with 1000 bp seeming to produce the most robust expression (Supplemental Figure 1B).

The integration and expression plasmid was assembled from a pRS426 backbone, bearing 1000 bp flanking sequences used for homologous recombination to the neutral *csr-1* locus, 1000 bp of the NCU04502 promoter positioned at the 5’ end of the insertion, followed by the coding sequence for SON-1 and a 9 amino acid linker. The different FP coding regions could be easily inserted into the plasmid downstream of *son-1*+linker and upstream of the 3’ flank of the *csr-1* locus (Fig. 1A).

**Figure 1.**
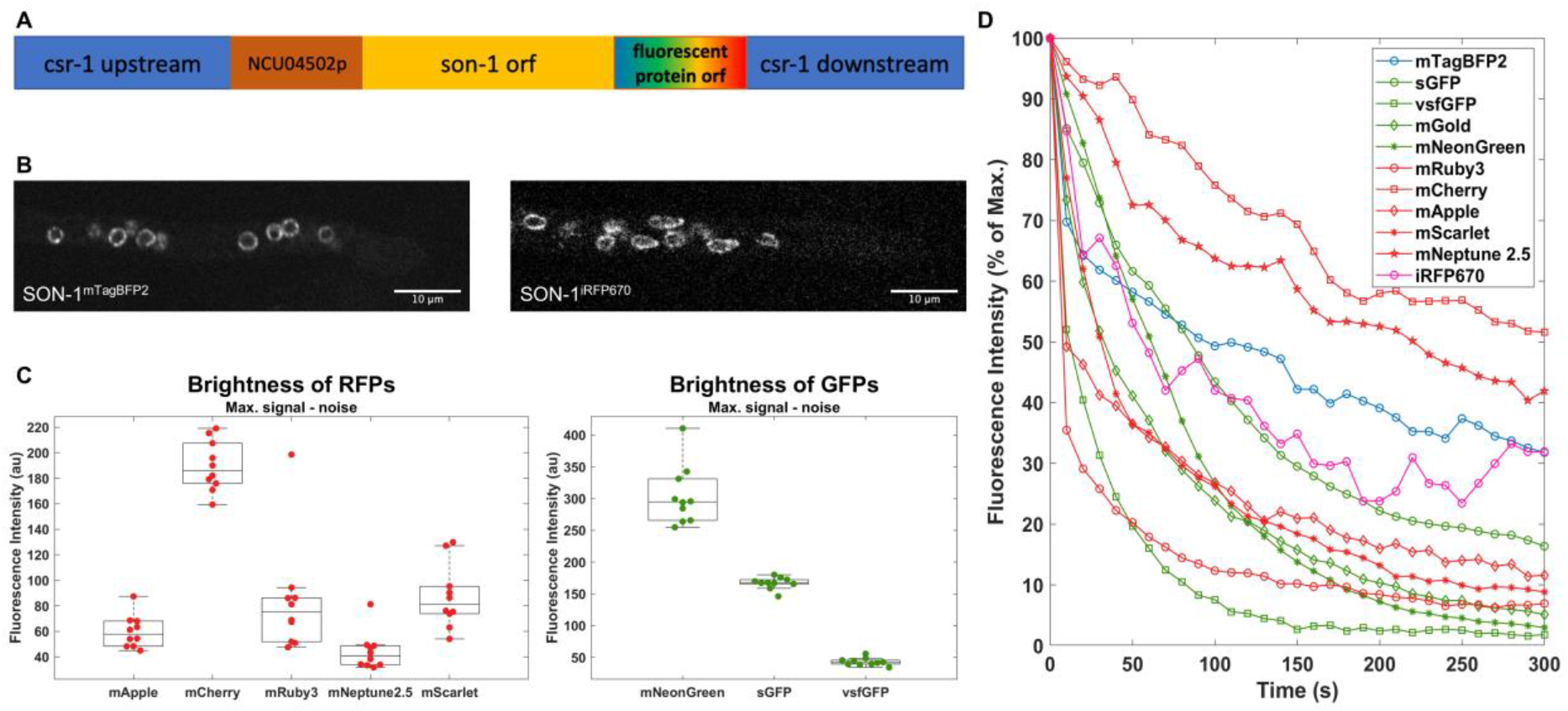
*In vivo* properties of constitutively expressed fluorescent proteins in *N. crassa*. A) Schematic of fluorescent proteins expressing genetic construct. 1 kb upstream and downstream targeting flanks for homologous recombination at the *csr-1* locus were inserted into pRS426. Original or codon-optimized open reading frames of fluorescent protein were inserted downstream of the NCU04502-promoter-driven SON-1 coding region into the vector using Gibson assembly. B) Confocal images showing mTagBFP2 or iRFP670 tagged SON-1 in hyphal tips. C) Fluorescent intensity measurements of RFPs and GFPs when tagging SON-1 in *N. crassa*. Each data point represents the mean of 10 measurements within 1 cell. For each fluorescent protein, n=10 cells. D) Bleaching profile of fluorescent proteins when tagging SON-1 in *N. crassa*. 95% of corresponding lasers were used for bleaching. Each curve represents the average of 3 individual measurements.

Green fluorescent proteins (GFPs) (Freitag et al., 2001) and red fluorescent proteins (RFPs) (Freitag and Selker, 2005) are the most commonly used fluorescent proteins in *N. crassa*. To extend the usable light spectrum for filamentous fungal cell biology research, we optimized the coding sequence of a rapidly-maturing monomeric blue fluorescent protein mTagBFP2 (Subach et al., 2011) to better fit the strong codon bias observed in *N. crassa* (Zhou et al., 2013). The optimized mTagBFP2 was successfully appended to SON-1 and produced bright *in vivo* signals (Fig. 1B). We also successfully visualized SON-1 constructs bearing a near-infrared fluorescent protein iRFP670 (Fig. 1B) and its monomeric version miRFP670-2 (data not shown). With these additional constitutively fluorescent proteins available, the potential of live-cell imaging in filamentous fungus has been largely improved.

We have also optimized various monomeric RFPs (mRFPs) and monomeric GFPs (mGFPs) not previously been widely used in filamentous fungi, tested their *in vivo* brightness and photostability properties, and compared them with the conventionally used monomers mCherry (Castro-Longoria et al., 2010) and sGFP (Freitag et al., 2001), respectively. mApple, mRuby, and mScarlet are among the best performing mRFPs and are brighter than mCherry in mammalian systems (Bajar et al., 2016b; Bindels et al., 2017; Shaner et al., 2004; Shaner et al., 2008). However, when appended to SON-1 in *N. crassa*, mCherry displayed the highest brightness and best photostability (Fig. 1C&D). We have also tested codon-optimized mNeptune2.5, which has a significantly more red-shifted emission spectrum than the other mRFPs while still being excited by a 561nm laser. This would allow more separation between the emission spectra during multi-color imaging and potentially reduce bleed-through. mNeonGreen is a very rapidly maturing bright monomeric yellow-green fluorescent protein derived from *Branchiostoma lanceolatum* (Shaner et al., 2013). Codon-optimized mNeonGreen produced significantly brighter signals than sGFP as expected (Fig. 1C), which makes it perfect for tagging proteins with low expression levels. Surprisingly, codon-optimized vsfGFP-9, which was predicted to have brightness similar to mNeonGreen, turned out to be very dim and sensitive to bleaching (Fig. 1C&D). These data confirm anecdotal observations suggesting that performance of fluorescent proteins in filamentous fungi is not always consistent with either theoretical predictions or measurements in mammalian systems.

### 2.2 Constitutive promoters driving diverse expression levels

In anticipation of needing a range of constitutive expression levels for different proteins, we tested a series of promoters using the same luciferase expression system as previously employed when characterizing the promoter of NCU04502, by inserting different promoters into the *csr-1::[promoter]-luciferase* plasmid. The promoters compared were *ccg-1* (*N. crassa*), *gpd* (*Cochliobolus heterostrophus*), NCU04502 (*N. crassa*), CMV immediate early, *ToxA* (*P. tritici-repentis*), and NCU08097 (*N. crassa*). The *ccg-1* promoter has long been a gold standard for a strong promoter in *Neurospora*. We described above the robust expression of the NCU04502 promoter, of which the 600 bp form was used for this project (Supplemental Table 1). The CMV immediate early promoter is widely used in animal systems as a strong promoter and is commonly found in mammalian expression vectors. The *gdp* and *ToxA* promoters were suggested by Michael Freitag (Oregon State University) as candidates for high and low expression, respectively. NCU08097 was identified as one of the lowest constitutively expressed genes in the Neurospora transcriptome with a log10 RPKM value of greater than 0 and less than 0.5 (Hurley et al., 2015). We constructed plasmids with 1500, 1000, and 500 bp of DNA upstream from the transcriptional start site and they all produced bioluminescence (data not shown), so we retained only the shortest variant for this comparison (Supplemental Table 1).

Transformation cassettes from the *csr::[promoter]-luc* plasmids were amplified by PCR and transformed into WT *N. crassa*. LUC expression was initially determined by placing mycelia of transformants growing on agar slants containing luciferin into a luminometer and measuring bioluminescence. Once the strains were established, they were analyzed by western blot using a luciferase antibody at 1:5000 (Abcam Cat. # ab185924), along with a tubulin antibody at 1:10000 (Sigma Cat. # T6199) as a loading control (Figure 2). Through this process, we identified promoter expression ranging from very strong to very weak. One surprising result was that the CMV promoter, which is commonly thought of as a very strong promoter in animals only provided a moderate level of expression in *Neurospora*.

**Figure 2.**
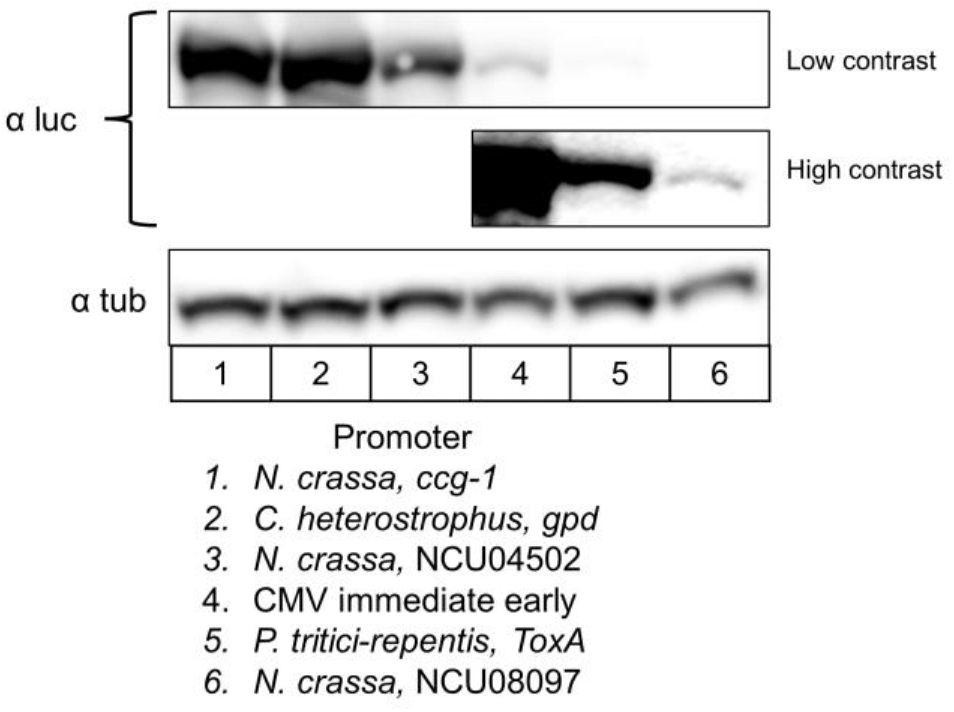
Analysis of promoters for varying levels of constitutive expression. Western blot analysis of luciferase to compare the relative strength of 6 different constitutive promoters for use in overexpression or heterologous protein constructs. 30 μg of total protein from *N. crassa* lysates were run in each well. Tubulin was used as a loading control. The upper and lower images of the luciferase blots were contrasted differently in ImageJ to make lane 6 visible. Lanes 1-3 were highly saturated at high contrast and therefore omitted from the second row.

### 2.3 4-color live cell imaging reveals 3D structure of *Neurospora crassa* mycelium

Incorporating multiple fluorescent protein tags into the same strain can be extremely useful to monitor the dynamic interactions between proteins and/or cell compartments. In recent years, 3 or 4-color live-cell imaging systems have been developed in mammalian cells and *S. cerevisiae* (Bajar et al., 2016a; Higuchi-Sanabria et al., 2016; Lee et al., 2013). However, to our knowledge no optimized fluorescent protein combination for 4-color imaging has been reported for filamentous fungi. Using the fluorescent protein tags we have adapted, we built a proof-of-principle strain using 4 different-colored fluorescent proteins to tag well-established cellular markers localized at different cellular locations.

We chose mTagBFP2 (blue), mNeonGreen (green), mApple (red), and iRFP670 (near Infrared) for which the spectra match with the classic excitation laser lines (405nm, 488nm, 561nm, 640nm) and CHROMA® Quad bandpass filter 89100bs, so that we can conduct fast 4-color imaging without switching between filters and minimizing bleed-through (Supplemental Figure 2). To prove that these tags do not interfere with correct localization, we tagged BML (β-tubulin) marking microtubules with mTagBFP2 (Freitag et al., 2004; Mouriño-Pérez et al., 2006), Septin CDC-11 marking septa, branching sites, and sub-apical compartments with mNeonGreen (Berepiki and Read, 2013; Riquelme and Martínez-Núñez, 2016), a core photoreceptor and clock protein WC-2 marking nuclei with mApple (Cheng et al., 2001; Schafmeier et al., 2008; Schwerdtfeger and Linden, 2000), and the nuclear pore complex (NPC) component SON-1 with iRFP670 (Roca et al., 2010). All fluorescent tags were added to the C terminus of the corresponding proteins. *cdc-11^mNeonGreen^* and *wc-2^mApple^* were edited at their endogenous locus. *bml^mTagBFP2^* and son-1^iRFP670^ were driven by a constitutive bidirectional promoter *(toxA* and *gdp* promoters, respectively) which we developed for this project and placed at the *csr-1* locus (Fig. 3A).

**Figure 3.**
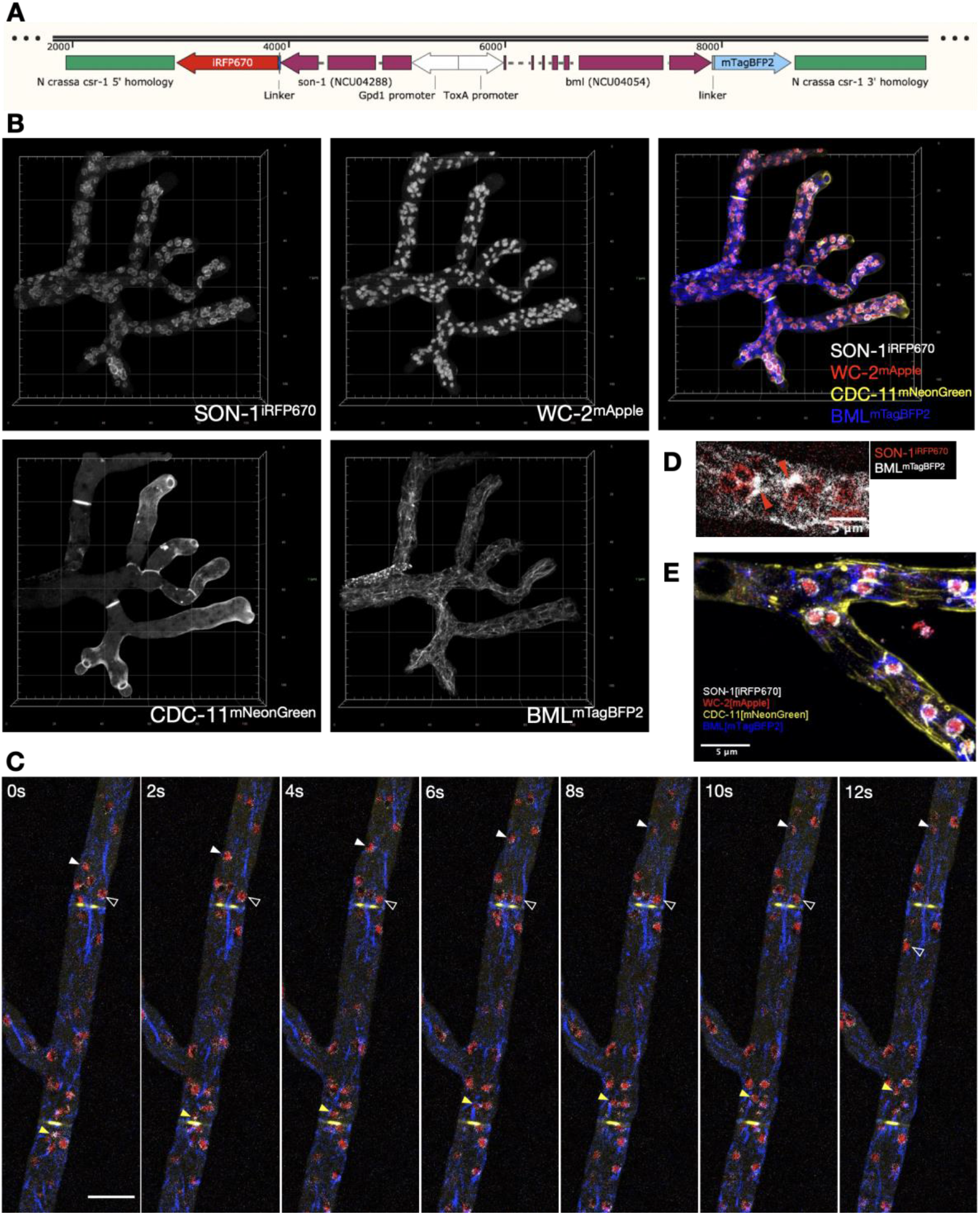
4-color live cell imaging reveals 4-dimensions cellular structures of *Neurospora crassa* mycelium. A) Schematic of BML and SON-1 tagging genetic construct using a bidirectional promoter. *C. heterostrophus gpd-1* promoter was used to drive the expression of *son-1* tagged by iRFP670. *P. tritici-repentis toxA* promoter was used to drive the expression of *bml* tagged by mTagBFP2. 1 kb upstream and downstream targeting flanks for homologous recombination at the csr-1 locus were inserted into pRS426. B) 3D rendered images of a hyphal tip from a culture grown on agar gel pads from the 4-color strain showing each channel or merged channel. C) Time lapse imaging of the living 4-color strain. The filled arrows (white and yellow) indicate nuclei moving along the cytoplasmic flow; the empty arrows indicate a nucleus moving against the cytoplasmic flow. Grey: SON-1^iRFP670^, Red: WC-2^mApple^, Yellow: CDC-11^mNeonGreen^, Blue: BML^mTagBFP2^. Scale bar = 10μm. D) Single focal image of the nuclear envelope (SON-1 in red) and microtubules (BML in greys). Red arrows indicate microtubule patches associated with the nuclear envelope. E) Z-projection of an unhealthy region in *N. crassa* mycelium network with ring-like and tubular septin structures (yellow).

Using confocal microscopy of a culture growing on agar gel pads (see Materials and Methods), we were able to capture structures of these cellular compartments in 4-dimensions in the same cell with high spatial resolution. All markers showed expected localization patterns (Fig. 3B & Supplemental movie 1). With 4-color live-cell imaging, we directly visualized the dynamic relationship between the cytoskeleton and nuclei inside the complex mycelium network. At the septum, microtubule bundles are squeezed through septal pores while retaining integrity. Nuclei were tethered to microtubules and traveled along the mycelium, through the septal pore, and went into different branches. Interestingly, besides the anticipated movements going with the cytoplasmic bulk flow (indicated by filled arrows), there were also nuclei dragged through the septal pore against the bulk flow (indicated by open arrows) in the same region (Fig. 3C & Supplemental movie 2). Microtubules form patches at the points where nuclei are tethered, and these patches are highly associated with the nuclear envelope (Fig. 3D). These observations suggest there may be a more direct interaction between microtubules and the nuclear envelope than current models describe (Mouriño-Pérez et al., 2016; Ramos-García et al., 2009). We have also observed a localization pattern of septins that has never been reported in *N. crassa*, in which in dying filaments, CDC-11 localizes to ring-like and tubular structures (Fig. 3D). A similar localization pattern of AspD, the CDC10 class of septins, has been reported in *Aspergillus fumigatus* hyphal tip (Juvvadi et al., 2011).

### 2.4 Photoconvertible fluorescent protein mEos3.1 helps monitor the dynamics of specific nuclei

Photoactivatable, photoconvertible and photoswitchable fluorescent proteins are relatively recent additions to the toolbox of live cell imaging research. They are known as “optical highlighters” due to their ability to change their excitation and emission spectrum upon stimulation. Photoconvertible proteins initially emit green light after excitation by a 488nm laser, but after stimulation (photoconversion) with UV light, they can then be excited by a 561nm laser instead and emit red light. Since the first report of a photoconvertible fluorescent protein in 2002 (Ando et al., 2002), such proteins have been adapted for a variety of uses. The first version of Eos was reported in 2004 (Wiedenmann et al., 2004), Dendra2 and mEosFP*thermo* were used in *Aspergillus nidulans* to monitor protein dynamics and to conduct super-resolution microscopy (Bergs et al., 2016; Zhou et al., 2018), and improved Eos variants continue to be developed. mEos3.1, a newer monomeric EosFP variant developed in 2012, displays higher brightness, faster maturation, and no incorrect localization caused by dimerization (Lippincott-Schwartz and Patterson, 2009; Zhang et al., 2012) but has never been adapted for use in filamentous fungi.

We codon-optimized the coding sequence of mEos3.1 for *N. crassa* and used it to tag histone H1 at its C terminus. We also added a V5 epitope tag between mEos3.1 and the open reading frame to allow confirmation of expression during troubleshooting and for the convenience of potential biochemical work. The expression of *hH1^mEos3.1_V5^* was driven by the *ccg-1* promoter and expressed from the *csr-1* locus (Fig. 4A), and this codon-optimized mEos3.1 fusion protein fluoresces properly. Before stimulation, hH1^mEos3.1^ could only be excited by a 488nm laser with no fluorescent signal showing up in the red channel. Using a 405nm laser we converted a subset of nuclei within a defined region of interest (ROI, shown as yellow squares) in the middle of a highly dynamic filament. After the illumination, hH1^mEos3.1^ within the exposed ROI were distinctly photoconverted to the red form and could be excited by 561nm laser (Fig. 4B). Converted mEos3.1 remained in its red-light emitting form, and thus could be used for long-term scanning (Hickey et al., 2004). We were able to track the movement of converted nuclei by monitoring the red channel (Supplemental Movie 3).

**Figure 4.**
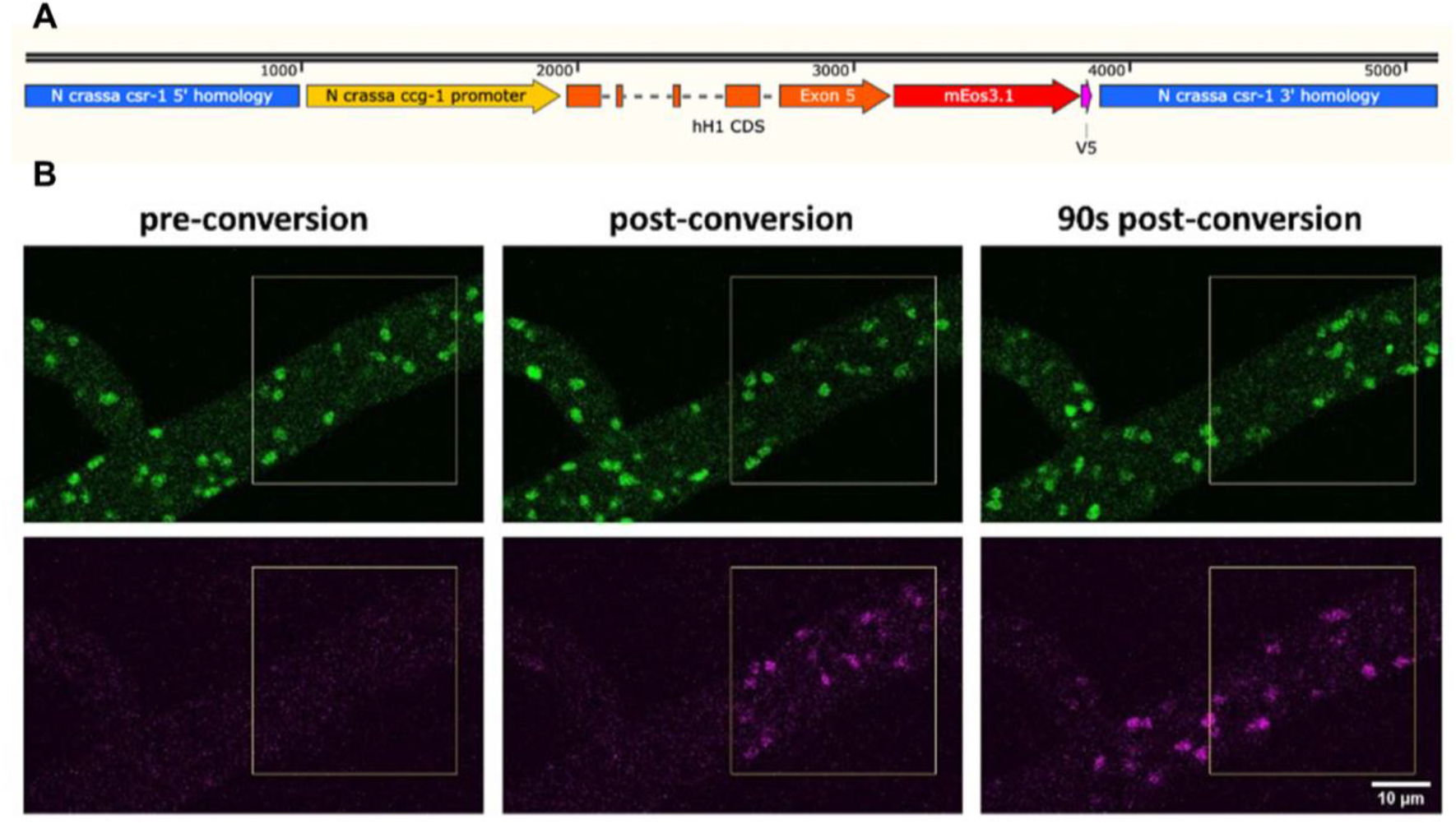
Tracking nuclear movement with a photoconvertible fluorescent protein mEos3.1. A) Schematic of *hH1*^mEos3.1^ tagging genetic construct. 1 kb upstream and downstream targeting flanks for homologous recombination at the *csr-1* locus were inserted into pRS426. The whole coding sequence of histone H1 followed by codon-optimized mEos3.1 and V5 epitope tag was inserted downstream of the *ccg-1* promoter into the vector using Gibson assembly. B) Time lapse imaging of histone H1 tagged with mEos3.1. The yellow square indicates the region chosen for photostimulation by 405 nm light to convert mEos3.1 from green to red. Images shown were acquired immediately before photoconversion, immediately after, and after 1.5 minutes of continuous imaging.

This proof-of-principal experiment showed the great potential of photoconvertible proteins. Besides monitoring traffic in a mycelial network, this tool can also be applied to other aspects of protein dynamic research (e.g. nucleocytoplasmic transportation, protein movements along the cytoskeleton, protein redistribution after fusion, etc.). In addition, mEos3.1 has properties suitable for recently developed super-resolution imaging techniques such as PALM and STORM.

## 3. Conclusions

The tools presented herein expand our ability to probe the cellular biology of *N. crassa* and should be applicable to a wide range of other filamentous fungi. The luciferase-based system for rapid testing of transcriptional promoters, set of promoters that express mRNA from very low to very high levels, and a customizable bidirectional promoter system can benefit researchers in a number of applications, not limited to fluorescence microscopy and cell biology. Analysis of brightness and bleaching among several fluorescent proteins in *N. crassa* provides valuable insights into which available fluorescent proteins might be best for various needs by revealing the tradeoffs one can expect surrounding brightness vs. bleaching when planning experiments. The ability to acquire images of four different proteins simultaneously facilitates design of experiments in which correlations and relationships of movements and activities among multiple independent proteins can be observed simultaneously. Finally, the optimization and characterization of a photoconvertible protein that permits a change in the color of proteins in discrete regions within a cell and for tracking their movement can promote a better understanding of how the fungal cytoplasm is organized, and how molecular trafficking works at a much deeper level.

## 4. Materials and Methods

### Plasmids and strains used

All plasmids reported in this work were constructed using NEBuilder® HiFi DNA Assembly Master Mix (New England Biolabs Cat. #E2621), based on the Gibson assembly method (Gibson et al., 2009). The common backbone plasmid used was pRS426, in which the multiple cloning site was replaced by the various DNA fragments described. The consensus wild type strain OR74A (FGSC#2489) was the parent strain for all experiments. In some cases, in which sexual crosses were required to obtain homokaryons, or when transforming into loci other than *csr-1, Δmus-51* strain were used.

### Western Blots

20 μg of total protein normalized by Bradford assay was run in each well of a pre-cast Bis-Tris gel (ThermoFisher Scientific, Catalog #NP0336) in MOPS buffer (ThermoFisher Scientific, Catalog #NP0001). The protein was transferred from the gel onto PVDF membrane using an iBlot gel transfer system (ThermoFisher Scientific). Antibody incubations were carried out in 5% nonfat milk in TBS buffer containing 0.2% Tween-20. Luciferase was detected using luciferase antibody (C-12) (Santa Cruz, sc-74548) as a primary antibody and HRP conjugated goat anti-mouse IgG secondary antibody (BIO-RAD, Catalog # #1706516). Chemiluminescence was detected using film for figure S1 and an Azure c400 imaging system (Azure Biosystems) for figure 2.

### Promoters and codon optimized fluorescent proteins

Promoter sequences for *N. crassa ccg-1*, NCU04502, and NCU08097 were retrieved from FungiDB; *gpd* (*Cochliobolus heterostrophus*) *and ToxA* (*P. tritici-repentis*) from JGI *Mycocosm* (https://mycocosm.jgi.doe.gov/). Among the fluorescent proteins used, coding sequences for sGFP (Freitag et al., 2004), mApple, mRuby3, mScarlet, mCherry, iRFP670 were used without modification, mApple-C1 was a gift from Michael Davidson (Addgene plasmid # 54631 ; http://n2t.net/addgene:54631 ; RRID:Addgene_54631) (Kremers et al., 2009). pKanCMV-mRuby3-18aa-Tubulin was a gift from Michael Lin (Addgene plasmid # 74256 ; http://n2t.net/addgene:74256 ; RRID:Addgene_74256) (Bajar et al., 2016b). pmScarlet_C1 was a gift from Dorus Gadella and the CMV immediate-early promoter described herein was amplified from this plasmid, as well as the mScarlet encoding gene (Addgene plasmid # 85042 ; http://n2t.net/addgene:85042 ; RRID:Addgene_85042) (Bindels et al., 2017). The CMV immediate-early promoter described the pCRE-iRFP670 was a gift from Alan Mullen (Addgene plasmid # 82696 ; http://n2t.net/addgene:82696 ; RRID:Addgene_82696) (Daneshvar et al., 2016). mNeonGreen was originally provided in a plasmid by Allele Biotechnology (Shaner et al., 2013). mNeonGreen, vsfGFP-9 (Eshaghi et al., 2015), mNeptune2.5 (Chu et al., 2014), and mTagBFP2 (Subach et al., 2011) were codon optimized by applying the Neurospora codon bias settings in SnapGene. All plasmids containing the SON-1-Fluorescent Protein fusions and their sequence data will be deposited at Addgene (www.addgene.org/) (Supplemental Table 2 & 3).

### Sample preparation

*Neurospora crassa* strains were grown overnight on agar pads (1x Vogel’s, 2% glucose, 0.17% arginine, 50ng/ml biotin, 1.5% agar) in Petri plates at 25°C starting from fresh conidia. Vegetative hyphae at the edge of growing colonies were imaged using the inverted agar block method (Hickey et al., 2004).

### Brightness measurements and Photobleaching

Strains were imaged using a Nikon Eclipse Ti-E microscope with Yokogawa CSU-W1 spinning disk system, Photometrics Prime BSI sCMOS camera, and piezo Z-drive. A Nikon LU-N4 laser launch that includes 405 nm, 488 nm, 561 nm and 640 nm lasers was used for excitation. The objective we used was Nikon CFI Plan Apochromat Lambda D 60X Oil objective (numerical aperture (NA) = 1.42).

For brightness measurements, 8μm Z-stacks with 300nm step size were taken for each strain using either 50% 561nm with 300ms exposure time (for RFPs) or 30% 488nm laser with 100ms exposure (for GFPs). The fluorescent intensities of 10 8×8 pixel ROIs with bright SON-1 signal and 10 8×8 pixel ROIs with cytoplasmic noise from each region were measured in ImageJ. The mean of 10 measurements was calculated and the noise-subtracted average SON-1 signal of each region was plotted.

95% corresponding excitation laser was used to perform photobleaching. Images were taken every 10s (300ms exposure time) for 5min. A 125×125 pixel ROI was created for each tip (and background) and the brightness of this region was measured in ImageJ. For each strain, the bleaching profiles of 5 healthy tips were measured. Background-subtracted means were normalized to that of the first timepoint of each time course and plotted.

### 4-color imaging

The 4-color strain was cultured as described above using the inverted agar block method (Hickey et al., 2004), and imaged using the Spinning disk confocal microscope as described above in the “Brightness measurements and Photobleaching” section with a CHROMA Quad bandpass filter (89100bs). 4 channels were imaged sequentially. 10μm Z-stacks or 1s-interval time series were taken. 3D rendering was performed in Arivis Vison4D software. Max projection and time series were processed in ImageJ.

### Photoconversion

Images were captured using Zeiss LSM880 equipped with an Airyscan detector, using a Zeiss Plan-Apochromat 63x/1.4 Oil DIC M27 objective. 6 iterations of a 20% 405nm laser were used to perform photoconversion within a selected ROI. Time series were recorded at 10s intervals with both 488nm and 561nm channels. Images were processed in Zeiss Zen software and labeled in ImageJ.

## CRediT authorship contribution statement

**Ziyan Wang:** Conceptualization, Methodology, Software, Validation, Formal analysis, Investigation, Data curation, Writing – Original Draft, Writing – Review & Editing, Visualization. **Bradley M. Bartholomai:** Conceptualization, Methodology, Software, Validation, Formal analysis, Investigation, Data curation, Writing – Original Draft, Writing – Review & Editing, Visualization. **Jennifer J. Loros:** Conceptualization, Resources, Supervision, Project administration, Funding acquisition. **Jay C. Dunlap:** Conceptualization, Resources, Writing – Review & Editing, Supervision, Project administration, Funding acquisition

## Acknowledgements

The *gdp* and *ToxA* promoters were gifts from Michael Freitag (Oregon State University). This work was supported by grants from the National Institutes of Health to J.C.D. (R35GM118021) and J.J.L. (R35GM118022), as well as Dartmouth’s BioMT NIH NIGMS COBRE grant, P20-GM113132.

## Supplemental Materials

**Supplemental Figure 1.**
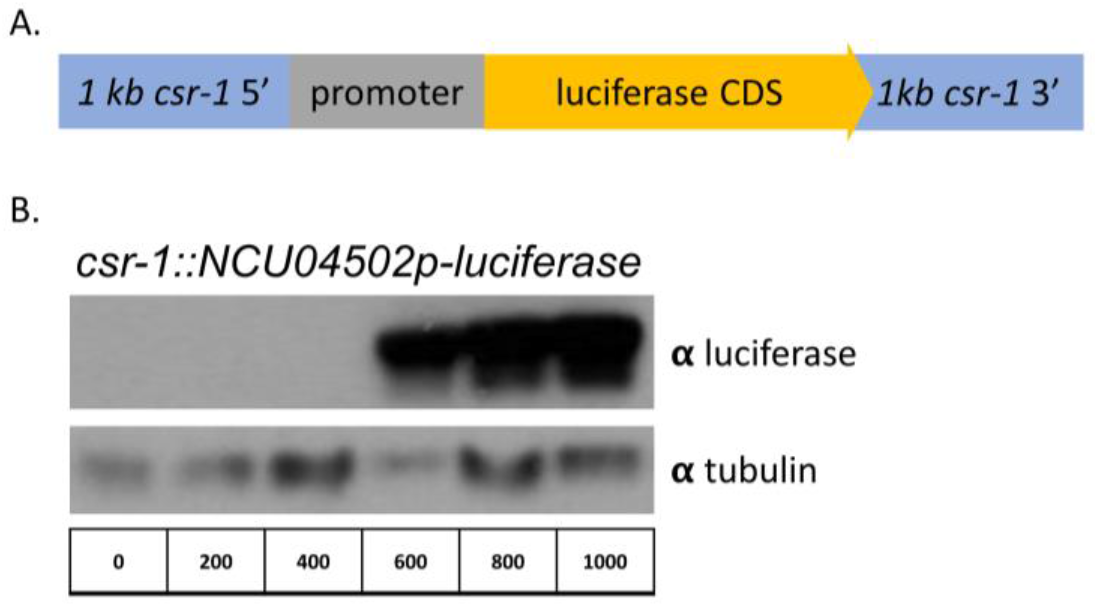
NCU4502 promoter length assay using a luciferase reporter. A) Schematic of luciferase reporter genetic construct. 1 kb upstream and downstream targeting flanks for homologous recombination at the *csr-1* locus were inserted into pRS426. Varying lengths of DNA up to 1 kb upstream of the translational start site (ATG) were inserted upstream of the firefly luciferase coding region into the vector using Gibson assembly. B) Western blot of luciferase expressed by varying lengths of NCU4502 promoter in liquid grown shaking culture under constant light at 25 °C.

**Supplemental Figure 2.**
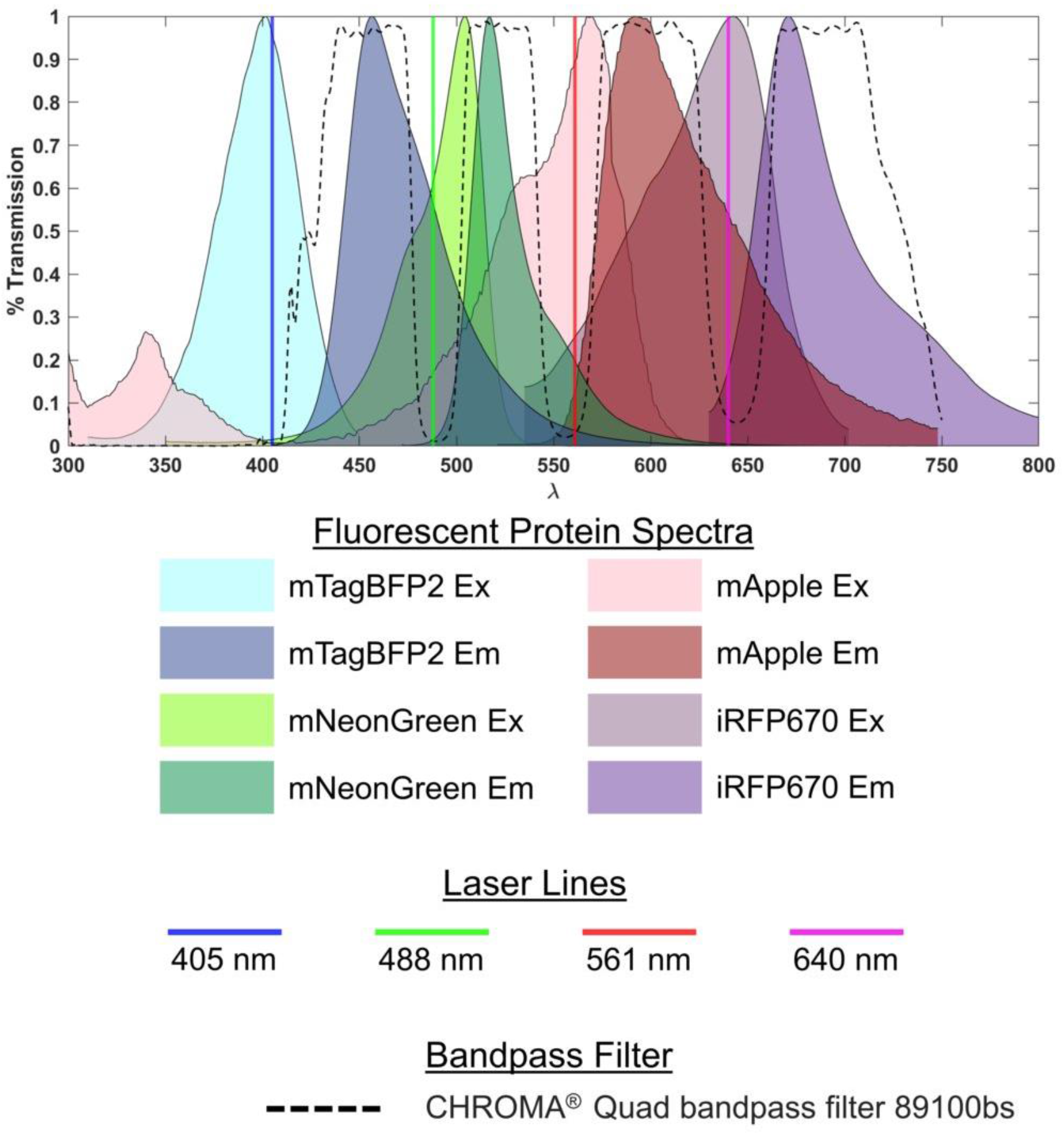
Fluorescence spectra, laser lines, and bandpass emission filters for 4 color imaging. Plot of the spectral properties of fluorescent proteins for use in 4-color imaging in *N. crassa*. Numerical data for spectra was acquired from FPbase.org and was replotted to create this figure. Laser lines shown are specific to the instrument the images were acquired on but are also common on many systems. Numerical data on band pass filters from Chroma (https://www.chroma.com/spectra-viewer).

**Supplemental Table 1.**
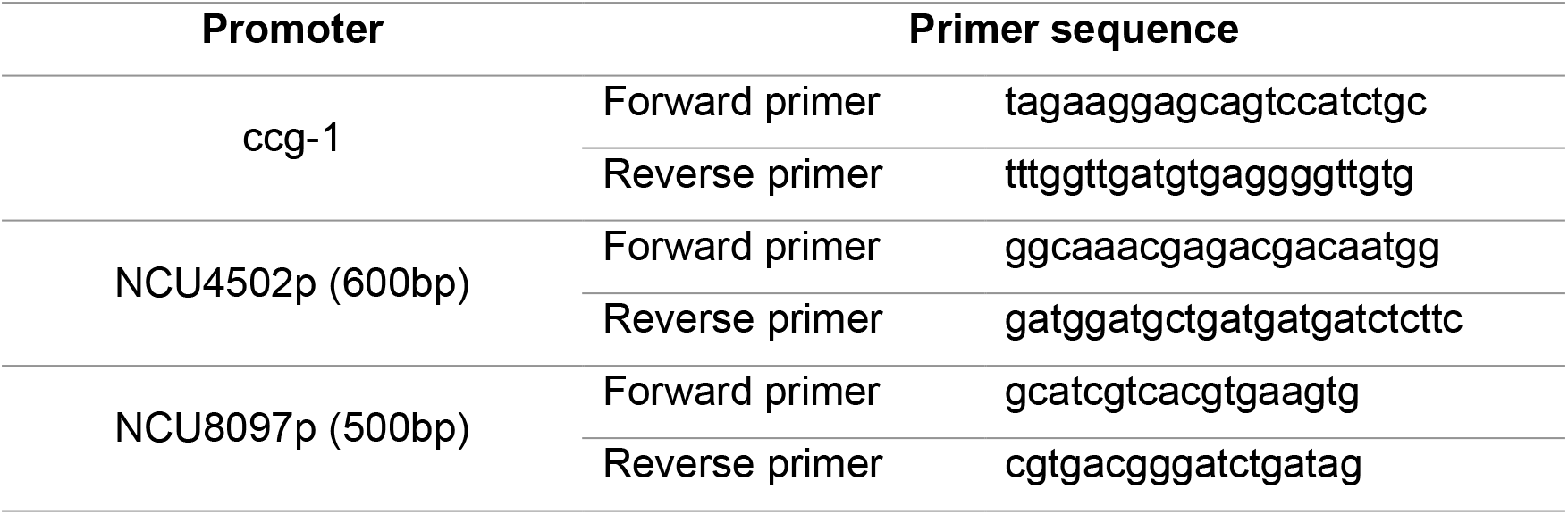
Primers used to clone out *N.crassa* promoters.

**Supplemental Table 2.** Plasmids used in this study.

| Plasmid | Lab ID | Addgene ID | Promoter | CDS |
| --- | --- | --- | --- | --- |
| csr-1_son-1_sGFP | pBB112 | 191561 | NCU4502p | son-1_sGFP |
| csr-1_son-1_mApple | pBB113 | 191741 | NCU4502p | son-1_mApple |
| csr-1_son-1_mCherry | pBB114 | 191742 | NCU4502p | son-1_mCherry |
| csr-1_son-1_mNeonGreen | pBB115 | 191743 | NCU4502p | son-1_mNeonGreen |
| csr-1_son-1_mRuby3 | pBB116 | 191744 | NCU4502p | son-1_mRuby3 |
| csr-1_son-1_mScarlet | pBB117 | 191745 | NCU4502p | son-1_mScarlet |
| csr-1_son-1_mTagBFP2 | pBB118 | 191746 | NCU4502p | son-1_mTagBFP2 |
| csr-1_son-1_vsfGFP9 | pBB119 | 191747 | NCU4502p | son-1_vsfGFP9 |
| csr-1_son-1_mNeptune2.5 | pBB120 | 191748 | NCU4502p | son-1_mNeptune2.5 |
| csr-1_son-1_iRFP670 | pBB122 | 191749 | NCU4502p | son-1_iRFP670 |
| csr-1_CMVp_luc | pBB507 |  | CMVp | luciferase |
| csr-1_gpdp_luc | pBB508 |  | gpdp | luciferase |
| csr-1_ccg-1p_luc | pBB509 |  | ccg-1p | luciferase |
| csr-1_ToxAp_luc | pBB510 |  | ToxAp | luciferase |
| csr-1_NCU4502p-600_luc | pBB511 |  | NCU4502p-600 | luciferase |
| csr-1_NCU8097p-500_luc | pBB514 |  | NCU8097p-500 | luciferase |
| csr-1_hH1_mEos3.1_V5 | pZW101 | 191750 | ccg-1p | hH1_mEos3.1_V5 |
| csr-1_iRFP670_son-1_GpdToxA_bml_mTagBFP2 | pZW102 | 191751 | Gpdp | son-1_iRFP670 |
|  |  |  | ToxAp | bml_mTagBFP2 |

**Supplemental Table 3.** *N.crassa* strains used in this study.

| Lab ID | Relevant Genotype | Figure(s) | Source/Reference |
| --- | --- | --- | --- |
| 2007 | *csr-1::NCU04502p_son-1<sup>mApple</sup>, mat A* | 1 | This Study |
| 2008 | *csr-1::NCU04502p_son-1<sup>mCherry</sup>, mat A* | 1 | This Study |
| 2009 | *csr-1::NCU04502p_son-1<sup>mRuby3</sup>, mat A* | 1 | This Study |
| 2010 | *csr-1::NCU04502p_son-1<sup>mScarlet</sup>, mat A* | 1 | This Study |
| 2011 | *csr-1::NCU04502p_son-1<sup>mNeptune2.5</sup>, mat A* | 1 | This Study |
| 2012 | *csr-1::NCU04502p_son-1<sup>mNeonGreen</sup>, mat A* | 1 | This Study |
| 2013 | *csr-1::NCU04502p_son-1<sup>vsfGFP-9</sup>, mat A* | 1 | This Study |
| 2014 | *csr-1::NCU04502p_son-1<sup>sGFP</sup>, ras-1<sup>bd</sup>, mat a* | 1 | This Study |
| 2015 | *csr-1::NCU04502p_son-1<sup>iRFP670</sup>, mat A* | 1 | This Study |
| 1661 | *csr-1::NCU04502p_son-1<sup>mTagBFP2</sup>, mat A* | 1 | This Study |
| 1965 | *csr-1::ccg-1p-luc, ras-1<sup>bd</sup>, mat a* | 2 | This Study |
| 1966 | *csr-1::CMVp-luc, ras-1<sup>bd</sup>, mat a* | 2 | This Study |
| 1967 | *csr-1::ToxAp-luc, ras-1<sup>bd</sup>, mat a* | 2 | This Study |
| 1968 | *csr-1::NCU04502p-luc, ras-1<sup>bd</sup>, mat a* | 2 | This Study |
| 1979 | *csr-1::NCU08097p-luc, ras-1<sup>bd</sup>, mat a* | 2 | This Study |
| 1980 | *csr-1::gdp-luc, ras-1<sup>bd</sup>, mat a* | 2 | This Study |
| 2016 | *wc-2<sup>mApple</sup>::hph+, cdc-11<sup>mNeonGreen</sup>::hph+,*<br>*csr-1::son-1<sup>iRFP670</sup>_gdpToxA_bml<sup>mTagBFP2</sup>,*<br>*ras-1<sup>bd</sup>, mat a* | 3 | This Study |
| 2061 | *csr-1::ccg-1p-hH1<sup>mEos3.1_V5</sup>, mat A* | 4 | This Study |

